# High-content imaging of live microglia-containing Alzheimer’s disease brain organoids

**DOI:** 10.64898/2026.10.26.761970

**Authors:** Daphne Quang, Rose Summers, Breanna R. Dooling, Hector Esquer, Qiong Zhou, Daniel V. LaBarbera, M. Natalia Vergara, Heidi J. Chial, Huntington Potter, Noah R. Johnson

## Abstract

Alzheimer’s disease (AD) is the most common progressive neurodegenerative disease for which there is currently no safe and effective treatment. AD is characterized by the pathological aggregation of amyloid-β (Aβ) in the brain resulting in neuronal dysfunction/death and cognitive decline. Modeling these processes *in vitro* remains challenging due to the lack of an immune-competent 3D *in vitro* human model of AD amenable to high-throughput analyses. Conventional cerebral organoids (COs) lack microglia and are optically dense which precludes real-time longitudinal analyses. Here, we employ the fluorescent amyloid probe PyrPeg to selectively monitor Aβ aggregation, and calcein AM and propidium iodide to assess cellular viability within microglia-containing COs (MCOs). We identify age-dependent DS-AD-associated phenotypes within T21 MCOs including Aβ neuropathology, neurodegeneration, and glial cell dysregulation. Our platform supports real-time, multiplexed tracking of disease progression and provides a scalable platform for longitudinal analyses studying disease mechanisms and targeted approaches to treatment.

**Motivation:** High-throughput *in vitro* drug screening using cerebral organoids (COs) has great potential to accelerate drug discovery efforts for treating Alzheimer’s disease (AD) and other neurodegenerative diseases. However, live fluorescence imaging of COs has proved challenging due to their dense, optically opaque tissue. Furthermore, conventional COs lack microglia which are key immunomodulators of pathogenesis in the AD brain. To address these limitations, we developed and optimized a live organoid imaging approach employing the fluorescent amyloid-binding probe PyrPeg to label amyloid-β (Aβ) aggregates within human induced pluripotent stem cell (hiPSC)-derived microglia-containing COs (MCOs). We achieved selective labeling and visualization of Aβ neuropathology in intact live MCOs and observed increased age-associated Aβ accumulation in MCOs derived from donors with trisomy 21 (T21) relative to disomy 21 (D21), consistent with the Down syndrome-associated AD (DS-AD) phenotype in the human brain. This method establishes a platform for high-throughput longitudinal drug screening in immunocompetent human COs that will enable the identification of new therapeutic candidates to modulate Aβ aggregation, glial reactivity, and neurodegeneration.

**Highlights:**

- A platform was developed for high-resolution *in vitro* visualization of insoluble Aβ aggregates in live MCOs.
- The approach enables deep tissue imaging of Aβ pathology in intact 3D MCOs, allowing spatial mapping of Aβ aggregate distribution over time.
- Differences in Aβ burden and viability of neurons, neural progenitors, and astrocytes were quantified, distinguishing T21 COs from D21 control MCOs.
- The platform is positioned to support high-throughput longitudinal drug screening, providing a scalable tool to identify novel therapeutics targeting Aβ-mediated neurodegeneration.

## Introduction

Alzheimer’s disease (AD) is the most common progressive neurodegenerative disease leading to cognitive impairment, progressive memory loss, and behavioral and psychological changes. AD is characterized neuropathologically by the accumulation of extracellular amyloid-beta (Aβ) plaques in the brain and vasculature which initiates a neuroinflammatory cascade, neurodegeneration, and ultimately cognitive decline^1,2^. Sequential cleavage of the amyloid precursor protein (APP) by secretase complexes generates the Aβ peptide, which then polymerizes into neurotoxic oligomers, fibrils, and ultimately parenchymal and vascular amyloid deposits as the earliest biochemical events leading to AD^3–5^. Because Aβ polymerization and amyloid deposition occur up to 20 years before the development of cognitive deficits, they are promising targets for therapeutic intervention. However, despite decades of research, translating these mechanistic insights into effective treatments has proved challenging, in part because the model systems currently available lack the ability to fully recapitulate disease pathology and are often not amenable to repeated longitudinal assessments. Advances in these two areas could ultimately enable the identification and development of effective treatments for AD.

The essential role of Aβ production in AD pathogenesis is supported by the discovery that autosomal dominant mutations in the γ-secretase components PSEN1 and PSEN2, or APP itself, are sufficient to cause early-onset AD^6–9^. Similarly, trisomy 21 (T21) in people with Down syndrome (DS) results in a third copy of the APP gene and is the most common form of genetically determined AD^10^. Indeed, all people with T21 develop DS-associated AD (DS-AD) brain amyloid neuropathology between the ages of 30-40, and 95% of those who live long enough develop clinical AD dementia within their lifetime^11,12^. As a result, AD is identified as a contributing cause of mortality for more than 70% of adults with DS beyond the age of 35^13^. Notably, a rare deletion of one copy of APP in some individuals with DS appears to avert DS-AD, underscoring the central role of APP gene dosage and Aβ accumulation in disease pathogenesis^14^.

Complexities of the human brain including cell type diversity, regional specificity, and heterogenous pathological features have made *in vitro* modeling of AD challenging^15,16^. However, the advent of techniques for the reprogramming of donor somatic cells to human induced pluripotent stem cells (hiPSCs) and their differentiation to neural cell types have enabled some models of cellular processes relevant to brain development and aging^17^. For example, hiPSCs can be used to generate cerebral organoids (COs) that recapitulate features of brain development in three-dimensional (3D) structures containing multiple subtypes of neuronal and glial cell populations^18,19^. COs exhibit greater cellular maturation and complexity than 2D monolayer cultures, and they exhibit physiological responses to aging-associated cellular stresses^20^. These features also enable the investigation of pathological mechanisms in human-derived brain models, which could accelerate the development and testing of new therapeutic strategies for AD and DS-AD. Indeed, COs derived from donors with AD exhibit some expected phenotypes including Aβ accumulation and amyloid plaque formation^19^.

Microglia play key roles in maintaining brain homeostasis and mediating immune responses in age-related neurodegenerative disorders including AD^21–23^. Efforts to model their roles have produced microglia-containing COs (MCOs), in which the cells appear to maintain their characteristic ramified morphology, inflammatory responsiveness, and phagocytic activities^24^. MCOs represent a considerable advance towards a physiologically relevant *in vitro* model capable of recapitulating the neuroimmune interactions that shape AD pathology^25^. In the human brain, glial cells are activated by neuritic plaques of insoluble Aβ, the abundance of which are strongly associated with synaptic loss and cognitive impairment^26–28^. Accurate quantitation of neuritic plaque formation and progression in models containing microglia and astrocytes are therefore essential to study the cellular and molecular mechanisms of AD pathogenesis.

Despite improvements in cellular diversity and tissue complexity, the usefulness of traditional CO systems is limited by their high opacity, making light-based measurements in real time difficult. Furthermore, conventional methods of immunolabeling neuritic plaques in fixed tissues are incompatible with high-throughput applications. To realize their full potential for mechanistic studies and therapeutic discovery, living organoids should be quantifiable by sensitive, real-time measurements of disease phenotypes. Fluorescent probes including Congo red, thioflavin-T, and Amylo-Glo, have enabled rapid, real-time quantitation of amyloid in live 2D models^29–31^; however, their short excitation wavelengths (< 500 nm) limit their use for 3D imaging due to shallow penetration depth (< 80 mm), autofluorescence, photobleaching, and phototoxicity^32^. As a result, near-infrared (NIR) amyloid probes including NIAD-4, CRANAD-2, and PyrPeg have been developed to overcome some of these drawbacks^33,34^. NIR probes have longer excitation wavelengths (> 700 nm), resulting in greater penetration depth and less light scattering in thick 3D tissues^35^. Moreover, NIR probes are compatible with high-content screening systems that enable automated, quantitative fluorescence imaging.

In this study, we establish a live organoid imaging platform that combines a physiologically relevant human-derived MCO model with high-content NIR optical imaging for real-time visualization of amyloid pathology. By integrating a human-relevant AD brain model with scalable quantitative imaging readouts, this platform provides a valuable tool for high-throughput screening of therapeutic candidates for AD and DS-AD.

## Results

### Characterization and validation of MCO models of DS-AD

We employed a protocol for generating MCOs^36^ and adapted it for the study of AD-related mechanisms. Two hiPSC lines were used: one derived from a male donor with T21 and one derived from an age- and sex-matched disomic control donor (D21). Neural progenitor cells (NPCs) and primitive macrophage progenitors (PMPs) were generated from the same hiPSC lines and combined at the embryoid body (EB) formation stage at a ratio of 7:3 NPCs:PMPs **(Fig. 1A)**, based on their physiological ratios in the human brain^37^. To confirm the presence of microglia, immunohistochemical (IHC) analyses of the microglial-specific markers Iba1 and P2RY12^37,38^ were performed in MCOs and in COs generated using the same number of NPCs but without PMPs included. In 90-d-old COs, we did not observe any cells expressing either Iba1 or P2RY12; however, in 90-d-old MCOs, we observed distinct microglia co-labeled for both Iba1 and P2ry12 **(Fig. 1B)**. Quantification confirmed that there were significantly more Iba1^+^ cells in the T21 MCOs than in the T21 COs **(Fig. 1C)**, representing approximately 10–15% of the total cells in MCOs, and quantification of P2RY12^+^ signal showed the same result (data not shown). Our findings are consistent with prior reports of microglia comprising approximately 8% of the total cells in MCOs^36^ and are also consistent with the 5–15% microglial composition of the human cortex^37^, thus validating our MCO development protocol.

**Figure 1.**
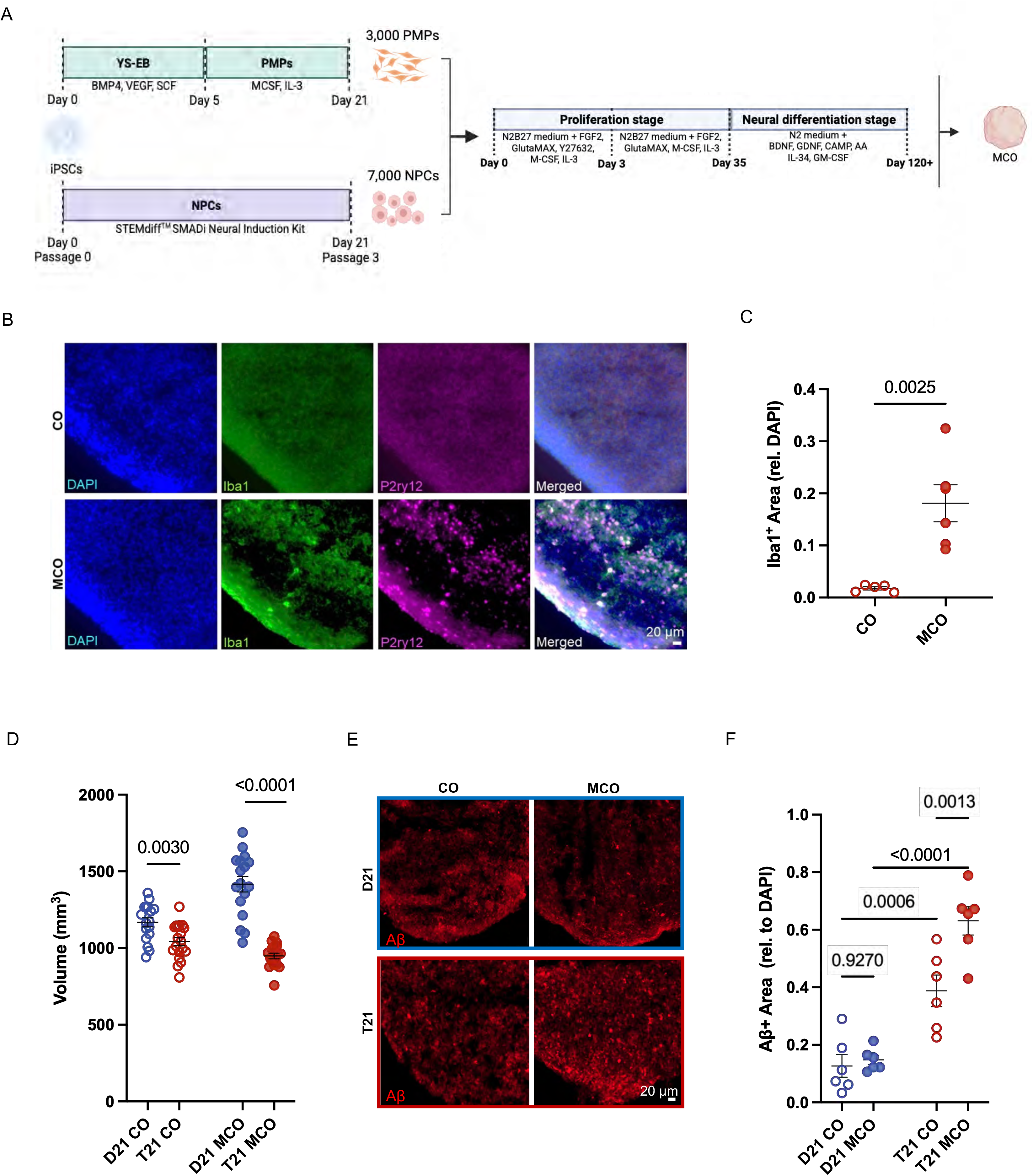
Microglia exacerbate AD neuropathology in T21 MCOs. (A) Schematic outlining the procedure for deriving MCOs by co-culture of iPSC-derived NPCs and PMPs under 3D conditions. (B) Representative immunohistochemistry images of sectioned 90-d-old T21 COs and MCOs stained for the microglia-specific markers Iba1 and P2RY12. All images were taken at 20x magnification. DAPI staining in blue. Iba1 was labeled with an Alexa 488-conjugated secondary antibody (green). P2ry12 was labeled with an Alexa 750-conjugated secondary antibody (purple). (C) Microglial quantification in T21 90-d-old COs and MCOs, measured by Iba1^+^ signal normalized relative to DAPI^+^ signal (p < 0.0025; unpaired t-test). The data represent the mean ± SD of n = 6 organoids per group. (D) Organoid volumes (mm^3^) in live, pre-fixed 120-d-old D21 and T21 COs and MCOs. The data represent the mean ± SD of n = 20 organoids per group. (E) Representative immunohistochemistry images of sectioned 120-d-old D21 and T21 COs and MCOs. All images were taken at 20x magnification. Aβ was labeled with an Alexa 555-conjugated secondary antibody. (F) Amyloid levels in 120-d-old T21 COs and MCOs in comparison to D21 COs and MCOs (p = 0.0006; p < 0.0001; two-way ANOVA) and T21 COs in comparison to T21 MCOs (p = 0.0013; two-way ANOVA) using the signal from Aβ labeled with an Alexa 555-conjugated secondary antibody normalized relative to the DAPI signal. The data represent the mean ± SD of n = 6 organoids, 4 slices/organoid for each group.

T21 COs have been shown to be smaller than D21 COs and to spontaneously develop AD neuropathology, including Aβ deposits^39^. To assess these phenotypes in our DS-AD MCO models, we collected 120-d-old organoids, measured their tissue volumes, and performed IHC analyses to detect Aβ. We found that T21 COs and MCOs had a significantly smaller volume than D21 COs and MCOs **(Fig. 1D)**. We observed numerous Aβ^+^ structures in T21 COs and MCOs, resembling intracellular Aβ aggregates and extracellular amyloid deposits **(Fig. 1E)**. Quantification revealed significantly more Aβ^+^ signal in T21 organoids relative to D21 organoids **(Fig. 1F**; F = 76.4400, *P* < 0.0001), and in MCOs relative to COs (F = 9.6457, *P* = 0.0056). Post-hoc analysis showed that the presence of microglia was associated with increased Aβ deposition in T21, but not D21, organoids **(Fig. 1F)**, indicating that microglia exert differential effects on Aβ depending on genotype. These data suggest that T21 genotype and the presence of microglia synergistically enhance Aβ deposition in DS-AD organoid models and validate their suitability for longitudinal studies of AD neuropathology.

### Optimization of an automated live imaging platform for DS-AD MCOs

We employed a spinning disk confocal-based high-content imaging system (Revvity Opera Phenix™ Plus) with a long pinhole-to-pinhole distance that effectively reduces out-of-focus noise to image whole MCOs in 96-well plates. To optimize the system for detection of amyloid in MCOs, we evaluated five different fluorescent live cell imaging probes: Amylo-Glo^31^, BioTracker NIR Amyloid-β Plaque Imaging Dye^40^, NIAD-4^41^, PyrPeg^34^, and ThT^30^ **(Fig. 2A)**. Additionally, 3D volume rendering of Z-stack images was performed to visualize acquired imaging depth **(Supp. Fig. S1)**. We found that all probes except NIAD-4 identified a significant increase in amyloid in 120-d-old T21 MCOs relative to D21 MCOs **(Fig. 2B)**; however, three probes (i.e., Amylo-Glo, BioTracker, and ThT) also exhibited relatively low signal intensity (< 2,500 a.u.). In contrast, PyrPeg produced markedly higher signal intensity (> 15,000 a.u.) and higher signal-to-noise ratio in T21 MCOs relative to D21 MCOs (**Fig. 2C**). These data suggest that PyrPeg provides greater dynamic range for detecting the effects of amyloid inhibitors in drug screening paradigms. Based on these metrics, we selected PyrPeg for further validation in our longitudinal live cell imaging experiments with MCOs.

**Figure 2.**
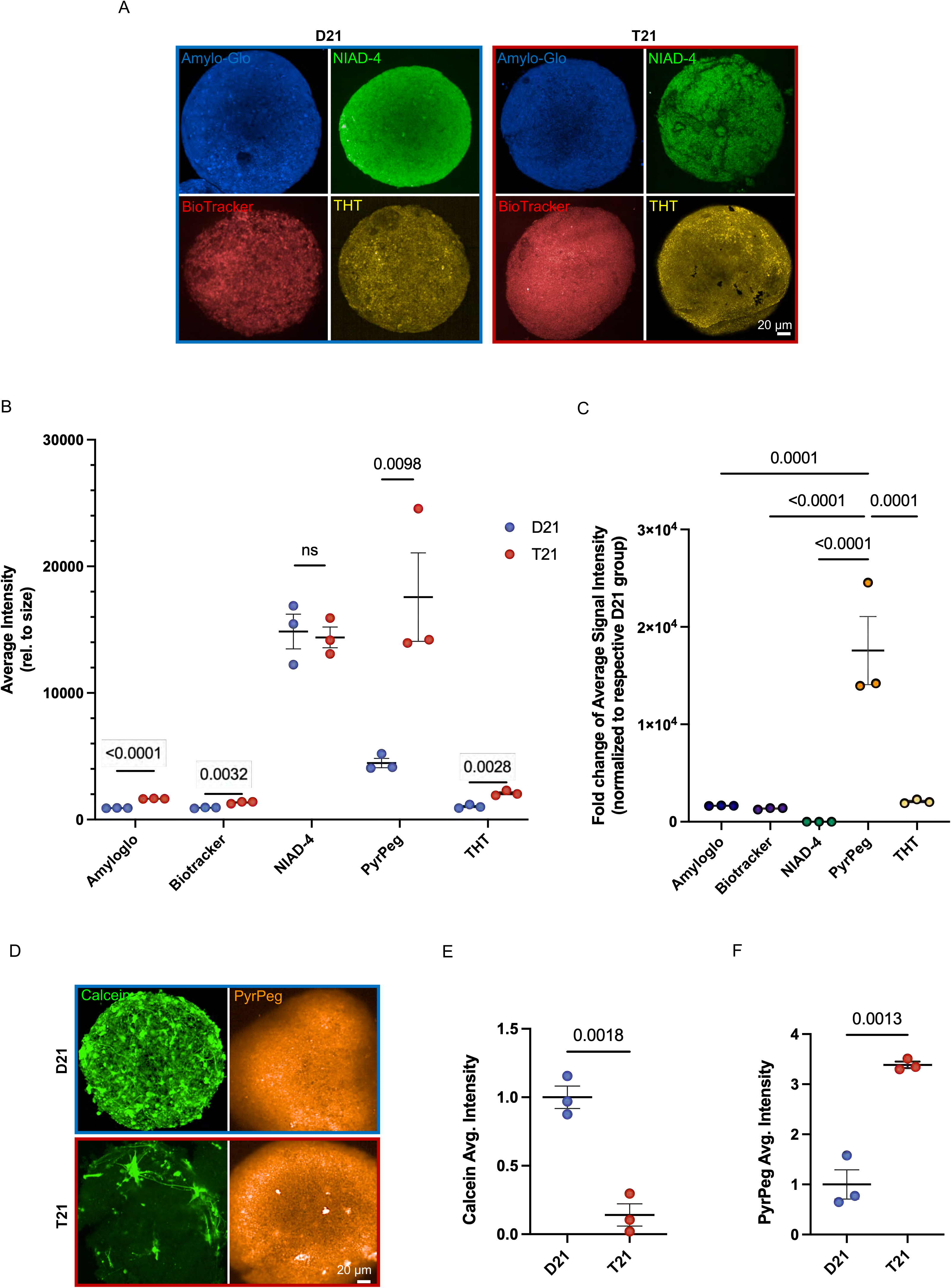
Opera Phenix™ Plus High-Content Screening system detects AD pathology in T21 MCOs. (A) Representative live cell images of intact 120-d-old D21 and T21 MCOs probed with various amyloid trackers. All images were taken at 20x magnification. Amylo-Glo (1x), labeled in blue; NIAD-4 (10 µM), labeled in green; BioTracker (10 µM), labeled in red; and THT (10 µM), labeled in yellow. (B) Comparison of amyloid trackers to detect Aβ in live, intact 120-d-old MCOs. All of the probes except for NIAD-4 showed a significant increase in Aβ in T21 MCOs. However, PyrPeg produced the highest signal intensity (~15,000 range; p = 0.0098; unpaired t-test). (C) Amyloid trackers normalized to the control group (D21 MCOs) to compare relative fold changes of average signal intensities across probes, with PyrPeg producing the highest fold change compared to all other probes (one-way ANOVA). (D) Representative live cell images of intact 120-d-old D21 and T21 MCOs showing decreased cell survival and increased amyloid in the T21 MCOs. All images were taken at 20x magnification. Live cell marker, calcein AM (1 µM), labeled in green. Aβ tracker, PyrPeg (10 µM), labeled in orange. (E) Quantification of cellular viability in 120-d-old T21 MCOs in comparison to D21 MCOs, using calcein AM average signal intensity of T21 MCOs normalized to D21 MCOs (p = 0.0018; unpaired t-test. (F) Quantification of Aβ signal in 120-d-old T21 MCOs in comparison to D21 MCOs, using PyrPeg average signal intensity of T21 MCOs normalized to D21 MCOs (p = 0.013; unpaired t-test). The data represent the mean ± SD of n = 3 organoids for each group.

To test the capacity of the platform for multiplexed analyses, we next evaluated calcein AM, a non-toxic dye that permeates intact cell membranes and fluoresces when hydrolyzed by intracellular esterases^42^. Furthermore, their distinct emission wavelengths allow calcein AM and PyrPeg to be multiplexed for future co-labeling in the same live MCOs (**Fig. 2D**). We observed substantially lower calcein AM fluorescence in T21 MCOs relative to D21 MCOs **(Fig. 2D)**, indicating reduced numbers of viable cells. Quantification revealed that T21 MCOs had a significant reduction in calcein AM^+^ signal **(Fig. 2E)** and a significant increase in PyrPeg^+^ signal relative to D21 MCOs **(Fig. 2F)**. These data indicate that greater amyloid deposition in DS-AD MCOs is accompanied by a reduction in cell viability.

### Selective vulnerability of neural cell types in DS-AD MCOs

To determine whether neurodegeneration underlay the reduced cell viability in T21 MCOs, we next performed IHC co-labeling of neuronal nuclei (NeuN) and caspase-3 (Cas-3), a key executioner protease in the apoptotic pathway^43^. We observed Cas-3^+^ cells and NeuN^+^ cells in both D21 and T21 MCOs **(Fig. 3A)**, yet there were significantly more Cas-3^+^ cells in T21 MCOs relative to D21 MCOs **(Fig. 3B)**, consistent with previous studies reporting increased Cas-3 protein expression in cells from individuals with DS^44^ and elevated procaspase-3 levels in AD brain tissue^45^, supporting a potential association between increased Cas-3 expression and disease-related cellular stress and apoptotic susceptibility. Co-localization analysis also revealed greater Cas-3^+^/NeuN^+^ signal in T21 MCOs relative to D21 MCOs **(Fig. 3C)**, indicating enhanced engagement of apoptotic machinery in T21 neurons. These data suggest that DS-AD MCOs exhibit both amyloidosis and neurodegeneration.

**Figure 3.**
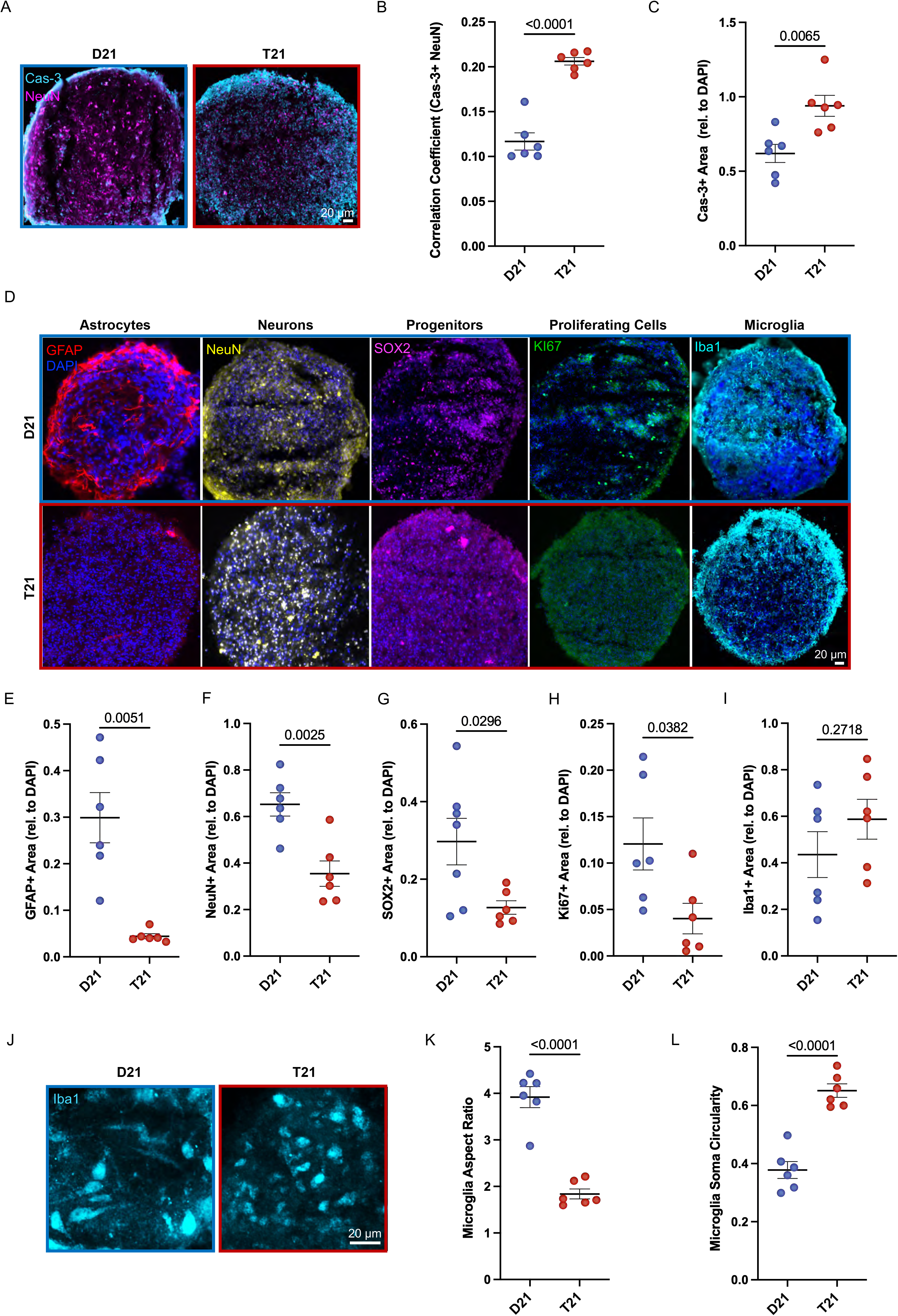
Cell-type specific alterations in T21 MCOs. (A) Representative immunohistochemistry images of sectioned 120-d-old D21 and T21 MCOs. All images were taken at 20x magnification. Caspase-3 (Cas-3) was labeled with an Alexa 647-conjugated secondary antibody. NeuN was labeled with an Alexa 750-conjugated secondary antibody. (B) Quantification of Cas-3 expression in 120-d-old T21 MCOs compared to D21 controls (p = 0.0065; unpaired t-test). (C) Comparison of colocalization between NeuN+ neurons and Cas-3+ cells in D21 and T21 MCOs, quantified using Pearson’s correlation coefficient (p < 0.0001; unpaired t-test). (D) Representative immunohistochemistry images of sectioned 120-d-old D21 and T21 MCOs. All images were taken at 20x magnification. GFAP was labeled with an Alexa 555-conjugated secondary antibody. NeuN was labeled with an Alexa 750-conjugated secondary antibody. SOX2 was labeled with an Alexa 647-conjugated secondary antibody. Ki67 was labeled with an Alexa 488-conjugated secondary antibody. Iba1 was labeled with an Alexa 750-conjugated secondary antibody. All images were taken from separate organoids. (E) Quantification of astrocytes (GFAP) in 120-d-old T21 MCOs compared to D21 controls (p = 0.0051; unpaired t-test). (F) Quantification of neurons (NeuN) in 120-d-old T21 MCOs compared to D21 controls (p = 0.0025; unpaired t-test). (G) Quantification of neural progenitors (SOX2) in 120-d-old T21 MCOs compared to D21 controls (p = 0.0296; unpaired t-test). (H) Quantification of proliferating cells (Ki67) in 120-d-old T21 MCOs compared to D21 controls (p = 0.0382; unpaired t-test). (I) Quantification of microglial populations (Iba1) in 120-d-old T21 MCOs compared to D21 controls (p = 0.2718; unpaired t-test). Data represent mean ± SD of n = 6 organoids, 4 slices/organoid for each group. (J) 40x magnification Z-stack images of Iba1 stained D21 and T21 MCOs from Fig. 3D. (K,L) Quantification of microglial aspect ratio (K) and soma circularity (L) in D21 and T21 MCOs. Data represent mean ± SD of n = 6 organoids, 10 cells/organoid for each group. p < 0.0001 for both comparisons.

We next tested whether certain neural cell types were selectively vulnerable to increased amyloid deposition or other DS-AD-related mechanisms in T21 MCOs. We performed IHC analyses for astrocytes (GFAP), neurons (NeuN), NPCs (SOX2), microglia (Iba1), and cell proliferation (Ki67) in 120-d-old MCOs **(Fig. 3D)**. We observed a significant reduction in astrocytes **(Fig. 3E)**, neurons **(Fig. 3F)**, NPCs **(Fig. 3G),** and Ki67^+^ proliferating cells **(Fig. 3H)** in T21 MCOs relative to D21 MCOs, but no difference in microglia **(Fig. 3I)**. However, morphological analysis of Iba1-positive microglia revealed significant differences between D21 and T21 microglia **(Fig. 3J)**. D21 microglia exhibited a significantly higher aspect ratio, indicating a more elongated morphology **(Fig. 3K)**, whereas T21 microglia displayed significantly higher soma circularity, consistent with a more rounded cell body **(Fig. 3L)**. Astrocytes appeared to be the most vulnerable cell type in T21 MCOs with an 85% reduction in total GFAP^+^ immunolabeling, which also appeared fractured and punctate, relative to the filamentous labeling observed in D21 MCOs (**Fig. 3D**). These data confirm that both neurons and astrocytes are vulnerable to cellular degeneration in T21 MCOs. The reduction in NPCs and cell proliferation suggest that T21 MCOs also exhibit impaired progenitor cell maintenance which may contribute to the observed differences in neuron and astrocyte populations.

### T21 MCOs exhibit age-dependent AD neuropathology

We next determined whether the AD neuropathological load increases over time in DS-AD MCOs, consistent with the age-associated progression in the human AD brain. We aged D21 and T21 MCOs beyond 120 days and used our live MCO imaging platform to assess amyloid burden with PyrPeg and the level of cellular apoptosis with propidium iodide (PI), a reliable marker of cell death^46–48^ **(Fig. 4A)**. We observed that 150-d-old T21 MCOs had significant increases in both PyrPeg^+^ **(Fig. 4B)** and PI^+^ signal, relative to 150-d-old D21 MCOs **(Fig. 4C)**. These results are consistent with our prior observations in 120-d-old MCOs and validate the use of PI in our live MCO imaging platform.

**Figure 4.**
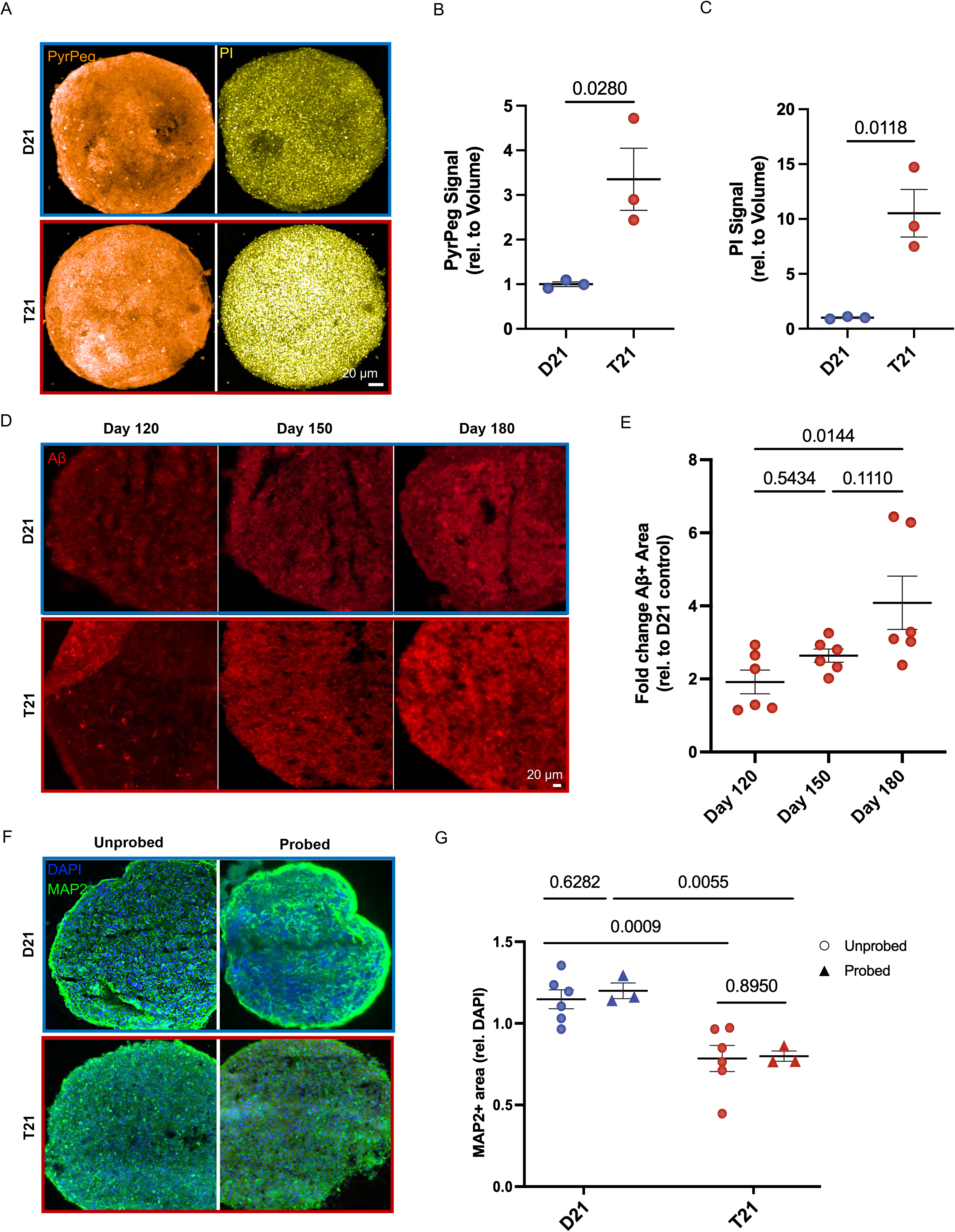
AD neuropathology increases in older T21 MCOs. (A) Representative maximum intensity projection images of live, intact 150-d-old D21 and T21 MCOs showing increased amyloid and increased cell death in the T21 condition. All images were taken at 20x magnification. Aβ tracker, PyrPeg (10 µM), labeled in orange. Dead cell marker, propidium iodide (PI, 1 µM), labeled in yellow. (B) Quantification of Aβ signal in 150-d-old T21 MCOs in comparison to D21controls using PyrPeg intensity normalized relative to organoid volume (p = 0.0280; unpaired t-test). The data represent the mean ± SD of n = 3 organoids. (C) Quantification of cell death in 150-d-old T21 MCOs in comparison to D21 control MCOs using PI intensity normalized relative to organoid volume (p = 0.0118; unpaired t-test). The data represent the mean ± SD of n = 3 organoids for each group. (D) Representative immunohistochemistry images of sectioned 120-, 150-, and 180-d-old D21 and T21 MCOs. All images were taken at 20x magnification. Aβ was labeled with an Alexa 555-conjugated secondary antibody. (E) Quantification of amyloid signal in 180-d-old T21 MCOs compared to 120-d-old T21 MCOs using the Aβ marker (p = 0.0144; one-way ANOVA). Data were normalized to the control group (D21 MCOs) to determine relative fold changes. The data represent the mean ± SD of n = 6 organoids and 4 slices/organoid for each group. (F) Representative immunohistochemistry images of sectioned 180-d-old D21 and T21 MCOs that never received dyes (unprobed) or that received dyes 30 days prior (probed). PyrPeg and PI were added once to 150-d-old MCOs, which were regularly fed until they reached 180 days of age. All images were taken at 20x magnification. DAPI staining is shown in blue. MAP2 was labeled with an Alexa 488-conjugated secondary antibody. Aβ was not assessed because PyrPeg and PI contributed to autofluorescence in their respective channels. (G) Quantification of unprobed and probed 180-d-old T21 MCOs in comparison to D21 controls, using the MAP2 neuronal marker normalized relative to the amount of DAPI signal (p = 0.0009; p = 0.0055; two-way ANOVA). Neuronal signal between unprobed and probed D21 and T21 MCOs was not significantly different (p = 0.6282; p = 0.8950; two-way ANOVA). The data represent the mean ± SD of n = 6 organoids for the unprobed groups and n = 3 organoids for the probed groups.

We next performed IHC analyses for Aβ in D21 and T21 MCOs at ages beyond 120 days **(Fig. 4D)**. We observed a significant effect of MCO age on Aβ deposition in T21 MCOs relative to D21 MCOs **(Fig. 4E; F = 5.4311, *P* = 0.0168)**, and post-hoc analysis confirmed a significant increase in Aβ in 180-d-old MCOs relative to 120-d-old MCOs **(Fig. 4E)**.

Finally, to evaluate the suitability of our live MCO imaging platform for longitudinal drug screening, we characterized the effects of PyrPeg and PI on neuronal viability by labeling 150-d-old MCOs with the probes and then performing IHC analyses 30 days later **(Fig. 4F)**. Quantification revealed that the MAP2^+^ signals from neurons in labeled MCOs, both D21 and T21, were identical to those of unlabeled MCOs **(Fig. 4G)**, indicating that the fluorescent probes do not compromise neuronal health or viability. Additionally, MAP2^+^ neurons were significantly reduced in 120-d-old T21 MCOs relative to D21 MCOs (F = 26.0, *P* = 0.0002), consistent with the neurodegeneration observed in younger T21 MCOs. Taken together, these data suggest that our live MCO imaging platform can precisely detect age-dependent changes in amyloid neuropathology and cellular health in T21 MCOs.

## Discussion

In this study, we address a key obstacle in AD research by developing a live organoid imaging platform that combines MCOs with fluorescent probes to monitor Aβ pathology and cellular health in real time. This system enables longitudinal, high-content imaging of insoluble Aβ aggregates within intact MCOs, providing functional readouts of disease pathology in a scalable, human-relevant format for drug testing and screening. Unlike conventional COs, which lack microglia, our MCO model includes PMPs along with NPCs, enabling self-organization of microglial populations within the 3D brain-like architecture. This is particularly relevant given that neuritic plaques *in vivo* are surrounded by activated microglia and astrocytes and elicit robust inflammatory responses, a key feature that is absent from most organoid models^49,50^. Microglia react to Aβ and influence its clearance, seeding, and toxicity, and they also activate other glial cells, highlighting the need for immune-competent model systems in AD^25,51^.

Our live organoid imaging approach was enabled by the use of PyrPeg, a NIR fluorescent probe that selectively binds to fibrillar Aβ and penetrates deep into 3D tissues with minimal phototoxicity and autofluorescence^34^. In contrast to conventional amyloid-binding dyes such as ThT and Congo red, PyrPeg offers high specificity for pathological Aβ fibrils, making it well-suited for quantification of amyloid in AD organoids^34,39^. When PyrPeg was multiplexed with calcein AM and PI for live/dead cell imaging, this platform revealed that T21 MCOs accumulate significantly more Aβ and undergo greater cell death compared to disomic controls, consistent with prior reports of DS-AD models^52–54^. Although calcein staining was limited in our T21 MCOs, it is relevant that calcein AM fluorescence reflects intracellular esterase activity and membrane integrity rather than viability alone. Cells undergoing significant stress or early stages of degeneration may exhibit reduced calcein uptake or retention prior to cell death^55^. Thus, the low calcein signal may reflect impaired cellular function associated with neurodegeneration rather than widespread irreversible cell death. Lastly, we found that PyrPeg exposure did not impair long-term cell viability, confirming the suitability of this approach for repetitive imaging in longitudinal drug screening applications.

T21 MCOs accumulated more Aβ than did T21 COs lacking microglia, emphasizing the importance of microglia-containing *in vitro* models for drug discovery. This system allows the evaluation of candidate drugs and therapies in a more physiologically relevant context than non-microglia containing counterparts, and can capture emergent neuroimmune interactions, such as inflammation, phagocytosis, and protein clearance. Beyond amyloid pathology, we identified cellular stress and survival as key AD-related pathways altered in T21 MCOs. Increased Cas-3 expression in T21 MCOs indicated greater activation/priming of apoptotic pathways and increased susceptibility to apoptotic signaling, particularly in neurons. T21 MCOs also had significant reductions in neurons, astrocytes, NPCs, and proliferating cells, suggesting impaired progenitor cell maintenance and reduced proliferative capacity. Collectively, these results indicate that T21 MCOs display enhanced cellular stress, disrupted cell maintenance, and features consistent with neurodegenerative processes relative to D21 MCOs.

In addition to these changes in cellular populations, morphological analysis revealed differences in microglia between the two groups. Microglia in T21 MCOs exhibited a more rounded morphology, characterized by a lower aspect ratio and higher soma circularity, compared with the more elongated, rod-shaped microglia observed in D21 MCOs. The morphological changes observed in microglia in T21 MCOs may reflect altered microglial states associated with disease, relevant to microglial alterations reported in the human DS-AD brain^56^.

Our results also indicate that T21 may exert an outsized effect on astrocyte populations in the DS brain. We observed a substantial reduction in GFAP^+^ astrocytes in T21 MCOs, consistent with reports characterizing glial cell dynamics in DS and DS-AD brains^57,58^. However, there are also contrasting reports of increased astrocyte populations and reactive astrogliosis during certain developmental windows and in specific brain regions^59–61^, suggesting that astrocytic population dynamics are context specific in the DS brain. Our findings indicate that T21 confers vulnerability not only to neurons, but also to astrocytes in an age-dependent manner, resulting in impaired maintenance of astrocyte populations. In COs, which lack systemic inflammatory or vascular cues promoting astrogliosis *in vivo*, this intrinsic susceptibility may be unmasked, highlighting a cell-autonomous deficit caused by T21. The resulting loss of astrocytes may exacerbate AD neuropathology by reducing neurotrophic support, altering extracellular homeostasis, or impacting metabolic and antioxidant functions. While astrocyte numbers decreased, we observed a trend towards increased microglia in aged T21 MCOs, potentially the result of microglial proliferation, a hallmark of many neurological diseases including AD^62^. This finding is particularly relevant in the context of DS-AD, where early and progressive alterations in glial populations and neuroinflammatory pathways occur in the human brain^63^. These observations suggest that our model recapitulates many aspects of DS-AD-associated glial dysregulation, providing a basis for further investigations of disease mechanisms and therapeutic interventions.

Coupling 3D amyloid imaging with simultaneous assessment of cell viability in live MCOs will enable mechanistic insights into the interplay between amyloid accumulation, glial dysfunction, and neurodegeneration. This platform provides a unique opportunity not only to evaluate Aβ-targeted therapeutics, but also to test interventions aimed at preserving cell type-specific functions. In doing so, our platform bridges the gap between 2D culture systems and *in vivo* models, offering a scalable and translationally relevant approach for accelerating therapeutic development in AD.

### Limitations of the study

Because microglia play a central role in Aβ clearance, neuroinflammation, and synaptic pruning^51,64,65^, their inclusion in our model system presents a key advantage over conventional COs; however, the phenotypic and transcriptomic states of the integrated microglia remain to be characterized, particularly in response to pharmacological agents. Future studies should incorporate transcriptional and phenotypic profiling of microglial states within MCOs at baseline and after pharmacological treatment. Single-cell transcriptomics and cytokine profiling could be used to delineate microglial phenotypes along disease and therapeutic trajectories. The live cell imaging techniques described here could be expanded in the future to include microglia-specific reporters (e.g., TMEM119-GFP) to enable live tracking of microglial migration and phagocytic activity in response to Aβ plaques^66^.

A second limitation of this study is the use of a single pair of hiPSC lines from age- and sex-matched D21 and T21 individuals. Genetic differences between the two donors, independent of chromosome 21, may have contributed to the observed phenotypes in MCOs. Future studies should include multiple pairs of matched hiPSC lines to better capture genetic variability and strengthen the generalizability of these findings. Alternatively, the use of isogenic hiPSC lines derived from the same donor mosaic for T21 could provide a rigorous approach to distinguishing the effects of T21 from those caused by genetic background.

## Resource availability

### Lead contact

Further information and requests for resources should be directed to and will be fulfilled by the lead contact, Dr. Noah Johnson.

### Materials availability

This study did not generate new unique reagents.

### Data and code availability

All data reported in this paper will be shared by the lead contact upon request.

## Acknowledgments

Research reported in this publication was supported by the National Institute on Aging of the National Institutes of Health (NIH) under award numbers R01AG078965 and RF1AG078965. This research was completed in collaboration with the Drug Discovery and Development Shared Resource at the University of Colorado Anschutz Medical Campus, RRID:SCR_021986, P30-CA046934, ALSAM Foundation, CU Anschutz Center for Drug Discovery.

## Declaration of interests

The authors have no competing interests to declare.

## Methods

### hiPSC Culture

The hiPSC D21 (LC68BC-5, male, 39) and T21 cell lines (LD49BC-3, male, 39) were obtained from the Gates Institute at the University of Colorado Anschutz. hiPSCs were plated onto Matrigel coated polystyrene 6-well tissue culture plates in mTeSR™ Plus Basal Medium supplemented with 1% Pen Strep and maintained at 37°C in a humidified 5% CO_2_ chamber. The medium was fully changed every day. The iPSCs were passaged every 4-6 days or until they reached approximately 70-80% confluency using ReLeSR™.

### MCO Generation

After hiPSCs were passaged twice, they were used to generate MCOs as described in a published protocol^30^. PMPs were differentiated as reported^30^; however, NPCs were differentiated using the STEMdiff™ SMADi Neural Induction Kit and corresponding protocol. A fixed ratio of PMPs and NPCs were then combined (specifically 7,000 NPCs and 3,000 PMPs) into each well of a 96-well ultra-low attachment (ULA) plate in MCO differentiation medium composed of 0.5X Neurobasal, 0.5X DMEM/F12, 1X N2, 1X B27-RA, 0.5X Glutamax, 10 ng/mL FGF-2, and 10 mM Y-2763 to form EBs, and were maintained at 37°C in a humidified 5% CO_2_ chamber. On day 3 following EB formation, half of the medium was replaced with fresh medium without Y-2763, and the MCOs were then fed every other day with MCO differentiation medium until day 35. From day 35 onward, the MCOs were fed every other day with MCO maturation medium composed of 0.5X Neurobasal, 0.5X DMEM/F12, 1X N2, 20 ng/mL BDNF, 20 ng/mL GDNF, 1 mM dibutyryl-cyclic AMP, 200 nM Ascorbic acid, 10 ng/mL GMCSF, and 100 ng/mL IL-34.

### Live-Cell Fluorescent Labeling of MCOs

All MCO medium was aspirated and replaced with fresh medium containing fluorescent probes prepared at the manufacturers’ recommended working concentrations. To facilitate probe penetration into the three-dimensional organoids, incubation times were doubled relative to those recommended for two-dimensional monocultures at 37°C under 5% CO2. The fluorescent probes used were Amylo-Glo (1× working concentration; 1h incubation; excitation/emission, 334/438 nm,), propidium iodide (2.5 µM; 1h incubation; 535/636 nm), NIAD-4 (10 µM; 1h incubation; 475/625 nm), Hoechst 33342 (2 µM; 30m incubation; 350/461nm), Thioflavin-T (10 µM; 1h incubation; 450/480 nm), Calcein AM (1 µM; 1h incubation; 500/521 nm), BioTracker (10 µM; 1h incubation; 500/700 nm), and PyrPeg (10 µM; 1h incubation; 444/598 nm). Following incubation, MCOs were washed three times with BrainPhys™ Imaging Optimized Medium before individual organoids were transferred to a new 96-well ultra-low attachment (ULA) plate containing 100 µL BrainPhys™ Imaging Optimized Medium for imaging.

Live cell imaging was performed by the University of Colorado Anschutz Drug Discovery and Development Shared Resource. Images were taken on the Revvity Opera Phenix™ Plus high content imaging system with Z-stacks at 20x. Wavelengths used were determined by manufacturers’ specifications for each probe.

Three separate fluorescence-based image analysis pipelines were executed using Acapella v5.6.0.132444 software, wherein each was tailored to the specific fluorophore channel for all probes. All analyses were performed on 3D image stacks with flatfield and brightfield corrections applied. For each dataset, the numbers of regions, as well as the means and standard deviations of the region volumes and mean channel intensities, were compiled for downstream statistical analyses.

### Histology and Immunostaining

At day 90, 120, 150, or 180 after EB formation, MCOs were rinsed once with Dulbecco’s Phosphate Buffered Saline (DPBS) and then fixed in cold 4% paraformaldehyde (PFA) in DPBS at 4°C for 15-20 h. After fixation, the 4% PFA was removed, and the MCOs were gently washed three times for 5 min each in DPBS which was then replaced with 30% sucrose solution with 0.01% sodium azide. MCOs were incubated in the sucrose solution at 4°C for 2-3 days. Afterwards, the MCOs were moved to TissueTek^®^ O.C.T. Compound in a cryostat mold and flash frozen. Frozen blocks were stored at −80°C. MCOs were sectioned on a cryostat at 20 μm on charged glass microscope slides. Slides were stored at −20°C.

For immunostaining, slides were washed in DPBS three times for 5 min each on an orbital shaker. DPBS was then replaced with a permeabilization/blocking solution (0.25% Triton X-100 in DPBS with 4% bovine serum albumin (BSA)) and put on a shaker for one hour. Slides were then inserted into a Sequenza™ rack and washed with DPBS once. Primary antibodies were added to the slides diluted in 0.1% Triton X-100 in DPBS with 4% BSA using the appropriate concentration for each antibody and incubated overnight at 4°C. Slides were then washed three times with DPBS. Secondary antibodies were diluted in 0.1% Triton X-100 in DPBS with 4% BSA and added to the slides, which were incubated in the dark for two hours at room temperature. The antibodies and concentrations used in this study are listed in the Key Resources Table. Slides were then washed three times with DPBS and the nuclei were then stained with Hoechst 33342 for 10 min at room temperature. Slides were then washed three times with DPBS and a coverslip was added along with Prolong™ Gold.

Images were acquired on an Olympus IX83 inverted fluorescence microscope. Images of organoids were captured at 20x magnification and then analyzed using Cell Sens v1.12 software (Olympus).

### Statistical analyses

Statistical analyses were performed using GraphPad Prism (version 10). For comparisons between two groups, two-tailed unpaired t-tests were used. For comparisons among three or more groups, one-way ANOVA was performed, followed by post hoc multiple-comparisons testing using the Holm-Šidák or Tukey method. For experiments involving two independent variables, ordinary two-way ANOVA was used to evaluate main effects and interactions. A P value < 0.05 was considered statistically significant, and all statistical tests were two-sided. Sample sizes, experimental replication, and the exact statistical tests used are detailed in the figure legends. Except in the case of MCO samples, where measurements were taken from multiple slices of the same MCOs, all measurements were obtained from distinct biological samples.

## Key resources table

| REAGENT or RESOURCE | SOURCE | IDENTIFIER |
| --- | --- | --- |
| **Primary and secondary antibodies** |  |  |
| Mouse Monoclonal anti-human anti-β-Amyloid (1:250) | BioLegend | 6E10 |
| Mouse Monoclonal anti-human GFAP (1:500) | EMD Millipore | MAB360 |
| Chicken Polyclonal anti-human IBA1 (1:1000) | Aves | IBA1-0020 |
| Chicken Polyclonal anti-human Ki67 (1:500) | Phosphosolutions | 1006-ki67 |
| Chicken Polyclonal | Phosphosolutions | 1100-MAP2 |
| anti-human MAP2 (1:1000) |  |  |
| Rabbit Polyclonal<br>anti-human NeuN (1:100) | abcam | ab177487 |
| Goat Polyclonal<br>anti-human Caspase-3 (1:1000) | Thermo Scientific | PA5142227 |
| Rabbit Polyclonal<br>anti-human P2RY12 (1:500) | Atlas Antibodies | HPA014518 |
| Goat Polyclonal<br>anti-human SOX2 (1:1000) | Mybiosource | MBS423840 |
| Donkey anti-Chicken IgY (H+L) Highly<br>Cross Adsorbed Secondary Antibody,<br>Alexa Fluor™ 488 (1:1000) | Thermo Scientific | A78948 |
| Donkey anti-Mouse IgG (H+L) Highly<br>Cross-Adsorbed Secondary Antibody,<br>Alexa Fluor 555 (1:1000) | Thermo Scientific | A-31570 |
| Donkey anti-Goat IgG (H+L) Highly<br>Cross-Adsorbed Secondary Antibody,<br>Alexa Fluor 647 (1:1000) | Thermo Scientific | A32849TR |
| Donkey Anti-Rabbit IgG Polyclonal<br>Antibody (CF™ 750) (1:1000) | Biotium | 20298 |
| HOESCHT 33432 (1.6 µm) | Thermo Scientific | 62249 |
| **Chemicals, peptides, and recombinant proteins** |  |  |
| DPBS | Gibco | 14190 |
| Growth Factor Reduced Basement<br>Membrane Matrix Matrigel | Corning | 356230 |
| mTeSR | STEMCELL Technologies | 100-0276 |
| Pen-Strep | Gibco | 15-140-122 |
| ReLeSR | STEMCELL Technologies | 100-0483 |
| Recombinant Human BMP-4 (HeLa<br>derived) | Peprtech | 120-05 |
| Recombinant Human SCF | Peprtech | 300-07 |
| Recombinant Human VEGF165 | Peprtech | 100-20 |
| X-VIVO™ 15 Serum-free Hematopoietic | Lonza | 02-053Q |
| Cell Medium |  |  |
| Recombinant Human M-CSF | Peprotech | 300-25 |
| Recombinant Human IL-3 | Peprotech | 200-03 |
| ACCUTASE™ | STEMCELL Technologies | 07920 |
| STEMdiff™ SMADi Neural Induction Kit | STEMCELL Technologies | 08581 |
| Neurobasal Medium | Gibco | 21103-049 |
| DMEM/F-12, HEPES | Gibco | 11330-032 |
| N-2 Supplement (100X) | Gibco | 17502048 |
| B-27 Supplement (50X), serum free | Gibco | 17504-044 |
| Recombinant Human FGF-basic | PeproTech | 100-18B |
| Y-27632 (Dihydrochloride) | STEMCELL Technologies | 72304 |
| Recombinant Human/Murine/Rat BDNF | PeproTech | 450-02 |
| Recombinant Human GDNF | PeproTech | 450-10 |
| Dibutryl-cAMP (Bucladesine) | Selleck Chemicals | S7858 |
| L-Ascorbic acid sodium salt | Thermo Scientific | 352680050 |
| Recombinant Human GM-CSF | PeproTech | 300-03 |
| Recombinant Human IL-34 | PeproTech | 200-34 |
| Amylo-Glo RTD | Biosensis | TR-300-AG |
| BioTracker NIR Amyloid-β Plaque Imaging Dye | Sigma-Aldrich | SCT073 |
| NIAD-4 | MedChem Express | HY-111263 |
| Pyrpeg Amyloid-B Dye | Sigma-Aldrich | SCT092 |
| Thioflavin-T | Biotium | 80033 |
| Calcein, AM | Thermo Scientific | C1430 |
| Propidium iodide | Sigma-Aldrich | P4170 |
| BrainPhys™ Imaging Optimized Medium | STEMCELL Technologies | 05796 |
| 16% Paraformaldehyde Aqueous Solution | Electron Microscopy Sciences | 15710 |
| D- Sucrose | Thermo Scientific | BP220-212 |
| 10% Sodium Azide | Thermo Scientific | S0209 |
| TissueTek O.C.T. Compound | Sakura Finetek USA Inc | 4583 |
| Triton X-100 | Sigma-Aldrich | 9036-19-5 |
| Bovine Serum Albumin (BSA) | EMD Millipore | 2930 |
| ProLong Gold Antifade Mountant | Thermo Scientific | P36934 |

---

**Supplementary Figure S1. Three-dimensional volume rendering of fluorescence in MCOs.** Representative Z-stacks were rendered in to visualize the spatial distribution of ThT fluorescence and acquired imaging depth of D21 and T21 MCOs.

**Supplemental Figure S2. Analysis sequences for live cell imaging of MCOs.** Diagram showing the analysis sequences used to determine signal intensity of the fluorescent live cell imaging probes Calcein, PyrPeg, and Propidium Iodide (PI) used to image MCOs with the Revvity Opera Phenix™ Plus high content imaging system.

## Notes

### Competing Interest Statement

The authors have declared no competing interest.

