## Supplementary figures and images for "High-content imaging of live microglia-containing Alzheimer’s disease brain organoids"

### Figure S1

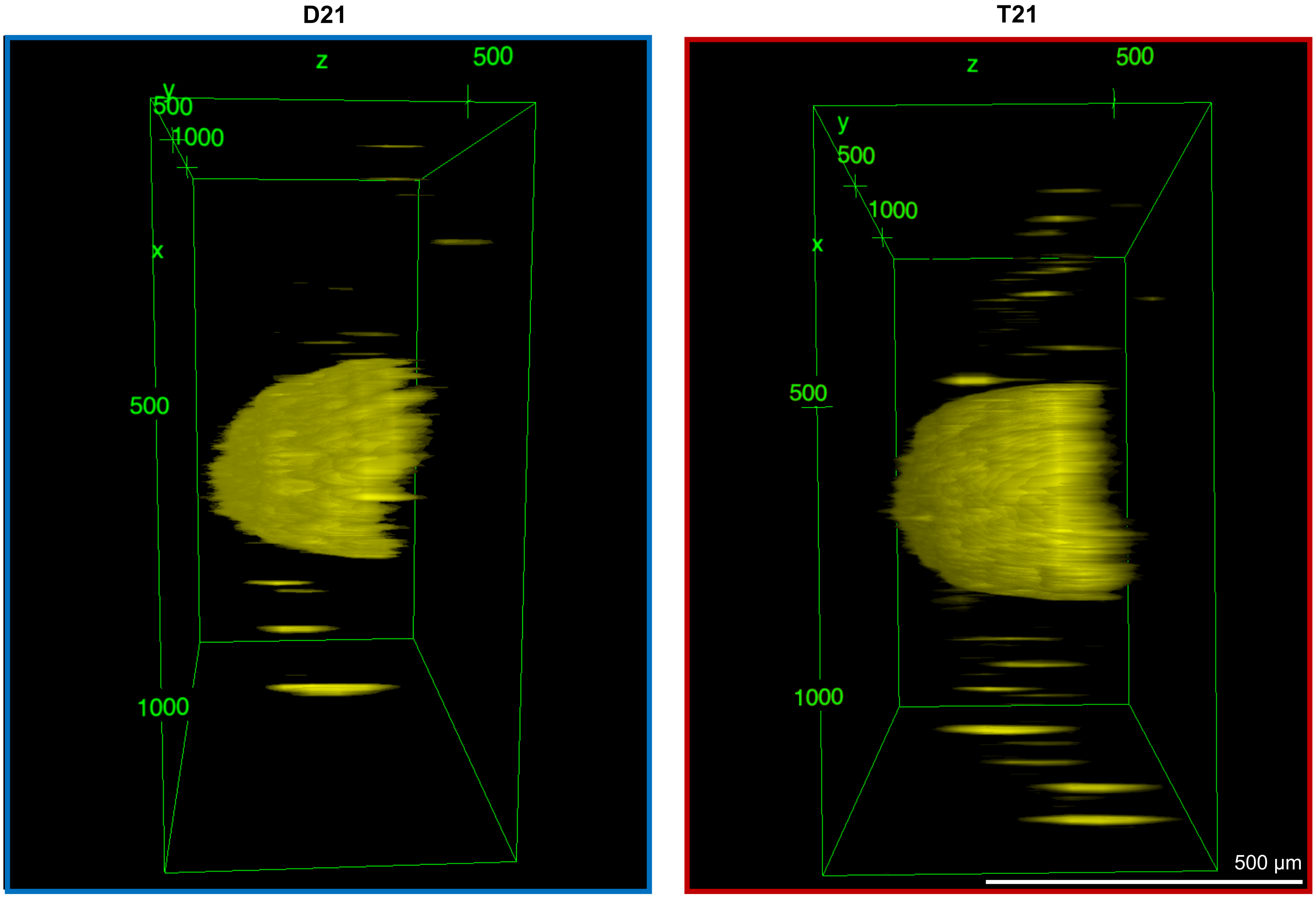


**Supplementary Figure 1**
