## Supplementary material for "High-content imaging of live microglia-containing Alzheimer’s disease brain organoids": Figure S2

### Analysis Sequence "Set 5 (Calcein Intensity)"

| Step | Input | Method | Output |
|---|---|---|---|
| **Input Image** | **Channel group :** 1<br>**Sequences :** ALL<br>**Flatfield Correction :** Basic Brightfield Correction<br>**Stack Processing :** 3D Analysis | | |
| **Find Image Region** | **Channel :** Calcein<br>**ROI :** None | **Method :** Local Threshold<br>Threshold : 0.4<br>Region Scale : 15 µm<br>Join touching Fragments<br>Volume : > 1000 $\mu m^3$ | Output Population : Image Region<br>Output Region : Image Region |
| **Calculate Morphology Properties** | **Population :** Image Region<br>**Region :** Image Region | **Method :** Standard<br>Volume | Property Prefix : Image Region |
| **Select Population** | **Population :** Image Region | **Method :** Filter by Property<br>Image Region Volume [$\mu m^3$] : > 100000<br>Image Region Volume [$\mu m^3$] : > 0<br>Boolean Operations : F1 and F2 | Output Population : Image Region Selected |
| **Calculate Intensity Properties** | **Channel :** Calcein<br>**Population :** Image Region Selected<br>**Region :** Image Region | **Method :** Standard<br>Mean | Property Prefix : Calcein Selected Intensity |

**Define Results** — Results

**Method :** List of Outputs
**Population : Image Region**
Number of Objects
Apply to All : None

**Population : Image Region Selected**
Number of Objects
Apply to All : Mean+StdDev
Image Region Volume [$\mu m^3$] : Mean+StdDev
Calcein Selected Intensity Mean : Mean+StdDev

**Object Results**
Population : Image Region : None
Population : Image Region Selected : None

### Analysis Sequence "Set 6 (PyrPeg Intensity)"

| Step | Input | Method | Output |
|---|---|---|---|
| **Input Image** | **Channel group :** 1<br>**Sequences :** ALL<br>**Flatfield Correction :** Basic Brightfield Correction<br>**Stack Processing :** 3D Analysis | | |
| **Find Image Region** | **Channel :** PyrPeg<br>**ROI :** None | **Method :** Local Threshold<br>Threshold : 0.4<br>Region Scale : 15 µm<br>Join touching Fragments<br>Volume : > 1000 $\mu m^3$ | Output Population : Image Region<br>Output Region : Image Region |
| **Calculate Morphology Properties** | **Population :** Image Region<br>**Region :** Image Region | **Method :** Standard<br>Volume | Property Prefix : Image Region |
| **Select Population** | **Population :** Image Region | **Method :** Filter by Property<br>Image Region Volume [$\mu m^3$] : > 100000<br>Image Region Volume [$\mu m^3$] : > 0<br>Boolean Operations : F1 and F2 | Output Population : Image Region Selected |
| **Calculate Intensity Properties** | **Channel :** PyrPeg<br>**Population :** Image Region Selected<br>**Region :** Image Region | **Method :** Standard<br>Mean | Property Prefix : PyrPeg Selected Intensity |

**Define Results** — Results

**Method :** List of Outputs
**Population : Image Region**

### Analysis Sequence "Set 2 (PI+PyrPeg)"

| Step | Input | Method | Output |
|---|---|---|---|
| **Input Image** | **Channel group :** 1<br>**Sequences :** ALL<br>**Flatfield Correction :** Basic Brightfield Correction<br>**Stack Processing :** 3D Analysis | | |
| **Find Image Region** | **Channel :** PyrPeg<br>**ROI :** None | **Method :** Local Threshold<br>Threshold : 0.4<br>Region Scale : 15 µm<br>Closing : 10 µm<br>Filling : Fill Plane-Wise<br>Join touching Fragments<br>Volume : > 10000 $\mu m^3$ | Output Population : Image Region<br>Output Region : Image Region |
| **Calculate Morphology Properties** | **Population :** Image Region<br>**Region :** Image Region | **Method :** Standard<br>Volume<br>Surface Area | Property Prefix : Image Region |
| **Select Population** | **Population :** Image Region | **Method :** Filter by Property<br>Image Region Volume [$\mu m^3$] : > 1000000<br>Image Region Surface Area [$\mu m^2$] : > 1000000<br>Boolean Operations : F1 and F2 | Output Population : Image Region Selected |
| **Calculate Intensity Properties** | **Channel :** Propidium Iodide<br>**Population :** Image Region Selected<br>**Region :** Image Region | **Method :** Standard<br>Mean | Property Prefix : Intensity Image Region Propidium Iodide |
| **Calculate Intensity Properties (2)** | **Channel :** PyrPeg<br>**Population :** Image Region Selected<br>**Region :** Image Region | **Method :** Standard<br>Mean | Property Prefix : Intensity Image Region PyrPeg |

**Define Results** — Results

**Method :** List of Outputs
**Population : Image Region Selected**
Number of Objects
Apply to All : Mean
Image Region Volume [$\mu m^3$] : Mean
Image Region Surface Area [$\mu m^2$] : Mean
Intensity Image Region Propidium Iodide Mean : Mean
Intensity Image Region PyrPeg Mean : Mean

**Method :** Formula Output
Formula : a/b
Population Type : Objects
Variable a : Image Region Selected - Image Region Volume [$\mu m^3$] Mean
Variable b : Image Region Selected - Intensity Image Region Propidium Iodide Mean Mean
Output Name : Volume/PI Intensity

**Method :** Formula Output
Formula : a/b
Population Type : Objects
Variable a : Image Region Selected - Image Region Volume [$\mu m^3$] Mean
Variable b : Image Region Selected - Intensity Image Region PyrPeg Mean Mean
Output Name : Volume/PyrPeg Intensity

**Object Results**
Population : Image Region : None
Population : Image Region Selected : None

**Supplementary Figure 2**
